# Unsupervised Neural Tracing in Densely Labeled Multispectral Brainbow Images

**DOI:** 10.1101/2020.06.07.138941

**Authors:** Bin Duan, Logan A Walker, Douglas H Roossien, Fred Y Shen, Dawen Cai, Yan Yan

## Abstract

Reconstructing neuron morphology is central to uncovering the complexity of the nervous system. That is because the morphology of a neuron essentially provides the physical constraints to its intrinsic electrophysiological properties and its connectivity. Recent advances in imaging technologies generated large quantities of high-resolution 3D images of neurons in the brain. Furthermore, the multispectral labeling technology, Brainbow permits unambiguous differentiation of neighboring neurons in a densely labeled brain, therefore enables for the first time the possibility of studying the connectivity between many neurons from a light microscopy image. However, lack of reliable automated neuron morphology reconstruction makes data analysis the bottleneck of extracting rich informatics in neuroscience. Supervoxel-based neuron segmentation methods have been proposed to solve this problem, however, the use of previous approaches has been impeded by the large numbers of errors which arise in the final segmentation. In this paper, we present a novel unsupervised approach to trace neurons from multispectral Brainbow images, which prevents segmentation errors and tracing continuity errors using two innovations. First, we formulate a Gaussian mixture model-based clustering strategy to improve the separation of segmented color channels that provides accurate skeletonization results for the following steps. Next, a skeleton graph approach is proposed to allow the identification and correction of discontinuities in the neuron tree topology. We find that these innovations allow our approach to outperform current state-of-the-art approaches, which results in more accurate neuron tracing as a tree representation close to human expert annotation.

## 1 Introduction

Brain circuits have long been known to arise from the physical connectivity patterns of many individual neurons (i.e., the “connectome”). It is valuable, therefore, for neuroscientists to describe the cable-like axon and dendrite morphologies to aid in the discovery of inputs and outputs of individual circuits and the roles of different cell types in these circuits (Zeng & Sanes, 2017). This presents a challenging paradigm for both data acquisition and analysis, as neural networks can encompass projection neurons spanning many centimeters and local connectivity features on the scale of hundreds of nanometers. These problems have begun to be addressed by ongoing technical innovations, however technical implementation details still require careful analysis.

Recent advances in light microscopy and genetic strategies for labeling defined groups of neurons have enabled neuroscientists to capture these dense volumetric images of neurons in the brain. Specifically, multispectral volumetric imaging of neurons, termed “Brainbow” (Figure 1**A**), has emerged as a promising approach to produce densely labeled brain samples (Livet et al., 2007; Cai et al., 2013). Briefly, individual neurons in a Brainbow sample each stochastically express combinations of fluorescent proteins, effectively labeling each neuron a different composite color. This enables unambiguous identification of individual axons and dendrites within a given volume. The continued improvement of Brainbow-like tools (Li et al., 2020), together with the advent of improved imaging technologies such as expansion microscopy (Chen et al., 2015; Tillberg et al., 2016) and light-sheet microscopy (Hillman et al., 2019; Jahr et al., 2015), makes Brainbow well-poised to make significant contributions to both connectomics and hypothesis-driven circuit analysis.

**Figure 1:**
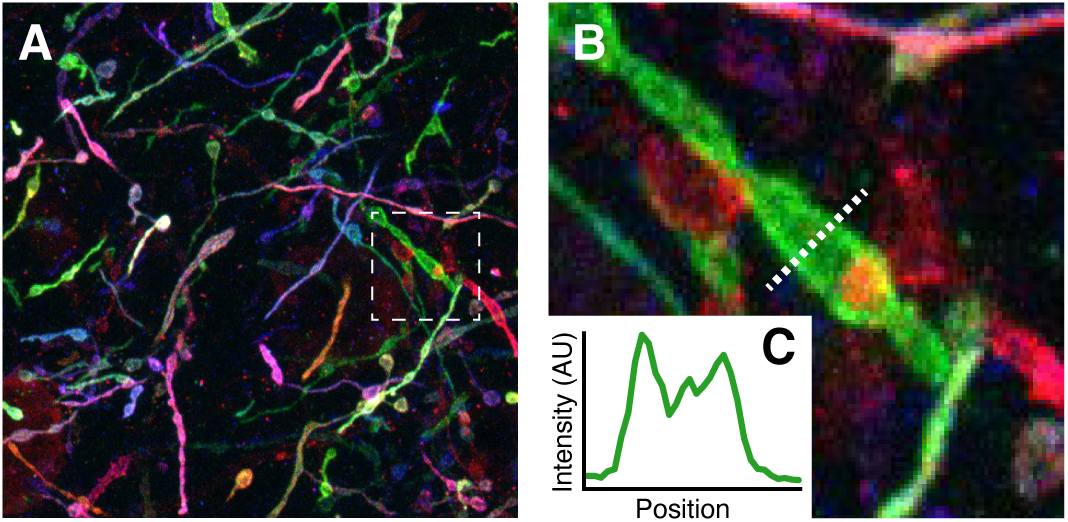
**A**, an example Brainbow image displaying the cable-like nature of axons and dendrites. **B**, a closeup of the box highlighted in **A**, displaying the hollow nature and noisy nature of Brainbow expression. **C**, a line intensity profile of the line in **B**. All images are 15-frame maximum intensity projections along the *z*-axis.

Despite the technological advances for collecting these micrographs, approaches for reliably analyzing and quantifying these rich datasets remain in their infancy. Along with large data sizes (in some cases as much as several terabytes), this is a difficult problem due to a large number of channels and imaging noise which is observed in the images (Figure 1**B–C**). Many tools exist for the manual or semi-manual reconstruction of neuron structures, such as *Neuromantic* (Myatt et al., 2012), commercial software such as *Neurolucida*. Recently, *nTracer* (Roossien et al., 2019) was released, specifically built for the analysis of Brainbow images. While these tools are effective for the reconstruction of small neuronal volumes, they are difficult to apply at scale due to the reliance on human input to the annotation, which can encompass tens to hundreds of human hours for even a relatively small image. Several automated solutions to aspects of the neuron segmentation problem have been proposed for use in both optical and electron microscopy generated images (Li et al., 2019; Gornet et al., 2019; Januszewski et al., 2018; Yang et al., 2019; Quan et al., 2016; Xiao & Peng, 2013), including several which specifically focus on Brainbow (Shao et al., 2012; Bas & Erdogmus, 2010; Wu et al., 2011; Sümbül et al., 2016). The earlier methods directly operate in voxel-level, which can be extremely computation-intensive and error-prone due to insufficient color consistency. Though the computation and color inconsistency issues are addressed in Sümbül et al. (2016), we can observe that this method results in fragmented (broken) neurite segmentations (Figure 2**D**, red arrows). These fragmented segmentations can be caused by a flaw in the supervoxelization process or occlusions by other neurons in the raw Brainbow data. In addition, none of the segmentation methods provide a tree-like neural tracing structure, which is required for further neuroinformatics analysis.

**Figure 2:**
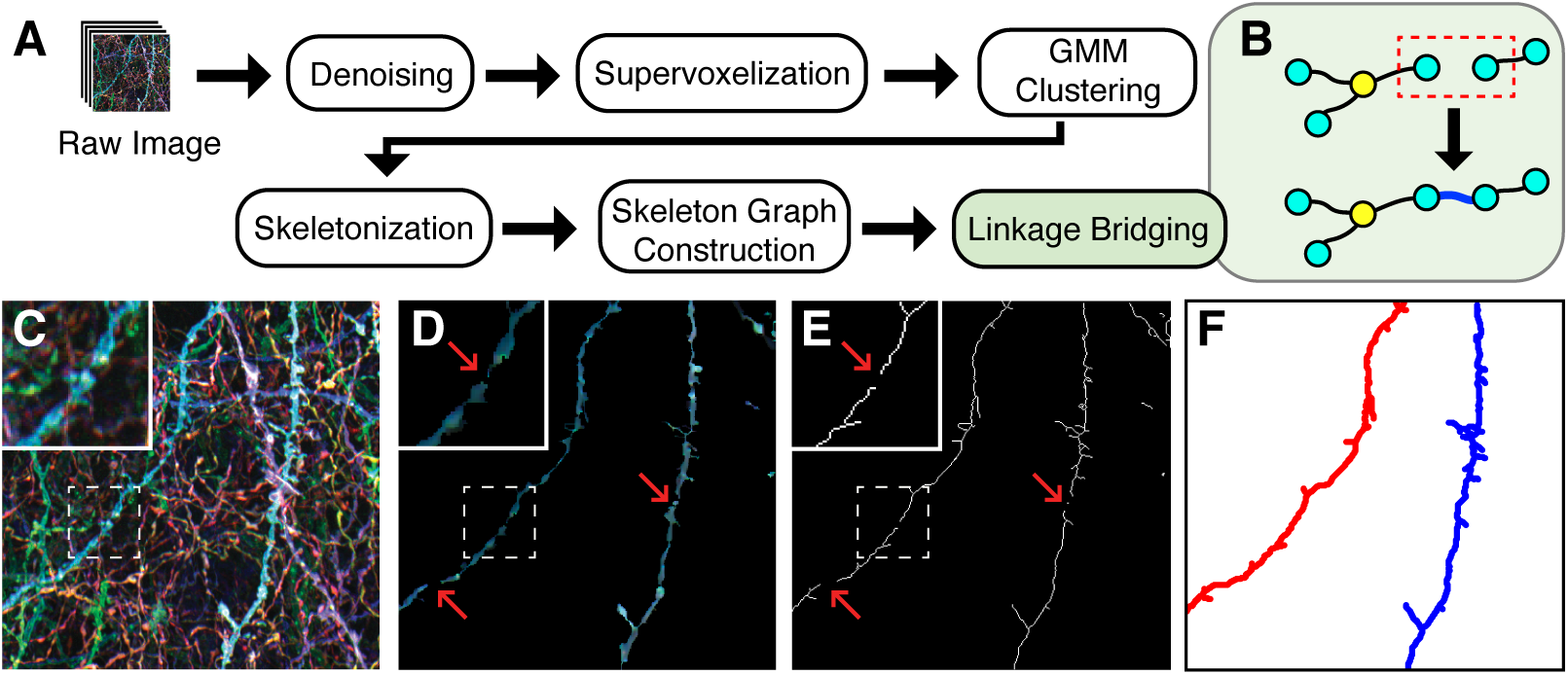
**A**, an overview of the data processing pipeline used in this paper. **B**, linkage bridging is used to repair broken trees, producing more robust neuron tracings. **C**, a raw Brainbow image. **D**, image **C** after GMM clustering is performed. **E**, image **D** after skeletonization is performed. **F**, the final reconstruction of the two neurites from **E**, pseudocolored for contrast. Arrows are used to identify places where there are erroneous breaks observed during skeletonization, which are corrected by linkage bridging. All images are maximum intensity projections of the entire image stack along the *z*-axis.

In this work, We intend to adapt the computationally efficient supervoxel-based segmentation used in Sümbül et al. (2016) but to address its fragmented segmentation problem. Aware of that the problem may be caused by its ke*rnel k*-means or spectral clustering, which are theoretically equivalent and tend to find circular or even-sized clusters (Dhillon et al., 2004), we instead use a probabilistic Gaussian mixture model. This modification allows the modulation of the distribution of supervoxel-representation to form more robust clusters and thus reduces the segmentation errors. Next, we extract a neuron tree topology (tracing), represented as a graph connectivity network, by skeletonization of the GMM-clustered segmentation. To address breaks in the neuron tracing which arise, we implement a graph-based method which utilizes the spatial relationship of the segmentation skeleton to bridge broken links, producing a more reliable tracing result.

Using this novel approach, we compare our automatic tracing to human-generated tracing, finding a high degree of agreement. Additionally, we show that the proposed approach can deal with loss of skeleton information and also more complicated neural structures within the input Brainbow images. In addition, whilst many hours can be spent for humans on reconstructing the neuroanatomy of such densely labeled multispectral Brainbow images, our proposed method can relieve the high demand for human annotation. This is a promising development, by which discovery of brain circuits can thus potentially be automated.

## 2 Methods

We first revisit the supervoxelization process, and then to solve the fragmentation problem, we make two innovations here: (i) To avoid biasing the segmentation being even-sized, we replace kernel *k*-means with Gaussian mixture model to modulate the supervoxel features which we obtain from the supervoxelization. (ii) To mitigate the gap between fragmented neurites, we develop a skeleton graph method (Figure 3) to reconstruct the tracing.

**Figure 3:**
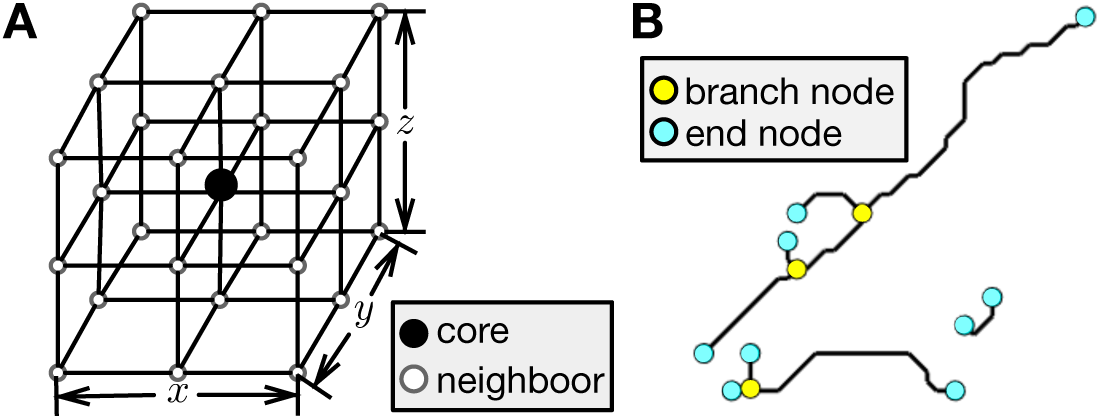
**A**, a diagram of 27-point stencil in euclidean space. **B**, example constructed graph visualization under *Definition 1*. Note that, the black lines between two nodes are following the path of the connected components using 27-point stencil.

### 2.1 Denoising

Images generated by confocal microscopy suffer from noise generated in both by stochastic variance in photon counts (shot noise) and from instrument noise (Sheppard et al., 1992). Figure 1**B** demonstrates a simple example of this, where the cross sectional intensity Figure 1**C** is found to be highly varied. To reduce these noise sources while preserving structures, we use a nonlocal transform-domain filter for volumetric data denoising. Assuming the noise is Gaussian, the BM4D denoiser (Maggioni et al., 2012) is applied in individual channels of the image stack, resulting in a denoised version.

### 2.2 Supervoxelization

Directly operating at the voxel level on Brainbow images is impractical due to computational complexity and erroneous due to the large observed color variances *in vivo*. One solution is to make supervoxels locally based on colors to oversegment the images (Sümbül et al., 2016). By making supervoxels, the computation is reduced, while also allowing more sophisticated features to be calculated at the supervoxel level. First, we restate two assumptions in a local and global context: (i) locally, neighboring voxels within the same neurites share similar colors; (ii) spatial information and color information act as a global constraint to obtain optimal segmentation. A watershed transform (Meyer, 1994) is used to generate supervoxels, and each supervoxel is summarized by its mean color in LUV space (Schanda, 2007).

After supervoxelization, we yield a set of supervoxels *S* = {*s*_*i*_}, where *s*_*i*_ denotes a supervoxel. The spatio-color constraint is imposed on supervoxel set *S*. We use a graph 𝒢 = (*V, E*) to represent the spatio-color relationship, where each supervoxel *s*_*i*_ in *S* represents a node in *V*. Edges are configured between pairs of supervoxels where: (i) the spatial distance between two supervoxels is less than *δ*_*s*_ or (ii) the color dissimilarity is less than *δ*_*c*_. In this way, the affinity matrix is constructed as

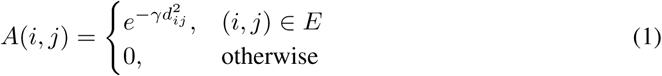

where *γ* is the Gaussian kernel parameter, and *d*_*ij*_ is the dissimilarity of color between two supervoxels. Note that the affinity matrix *A ∈*ℝ^*N ×N*^ is a sparse matrix where *N* represents the number of supervoxels. Instead of using *A* directly, we decompose A into lower dimension *d* by utilizing eigenvalue decomposition algorithm (Baglama et al., 2003), where *d* is the number of eigenvectors with the *d* largest eigenvalues. We denote the updated affinity matrix as our feature **X** *∈*ℝ^*N ×d*^ which is of the form

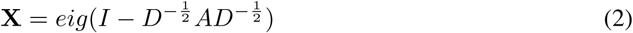

where *I* denotes identity matrix, and *D*_*ii*_ Σ = _*j*_ *A*_*ij*_, so that the eigendecomposition is imposed on the symmetric normalized Laplacian matrix.

### 2.3 Neuron Segmentation

Supervised learning methods (e.g., Gornet et al.) rely on a volumetric ground truth segmentation, which is difficult and, in many cases, infeasible. Thus, unsupervised approaches, such as kernel *k*-means and spectral clustering, allow more efficient solutions. However, after applying kernel *k*-means, we observe imperfect segmentation results, particularly with fragmentation near differences in neuron caliber (Figure 4). Thus, instead of imposing hard clusters on **X** via kernel *k*-means or its variants (Dhillon et al., 2004), we approximate the distribution of the feature **X** using mixture of Gaussians (GMM). We hypothesize that GMM will perform better because it does not bias the cluster sizes to have specific structures as does kernel *k*-means (Circular). We use the expectation-maximization (EM) algorithm to iteratively optimize the model and check the variational lower bound until convergence is accomplished.

**Figure 4:**
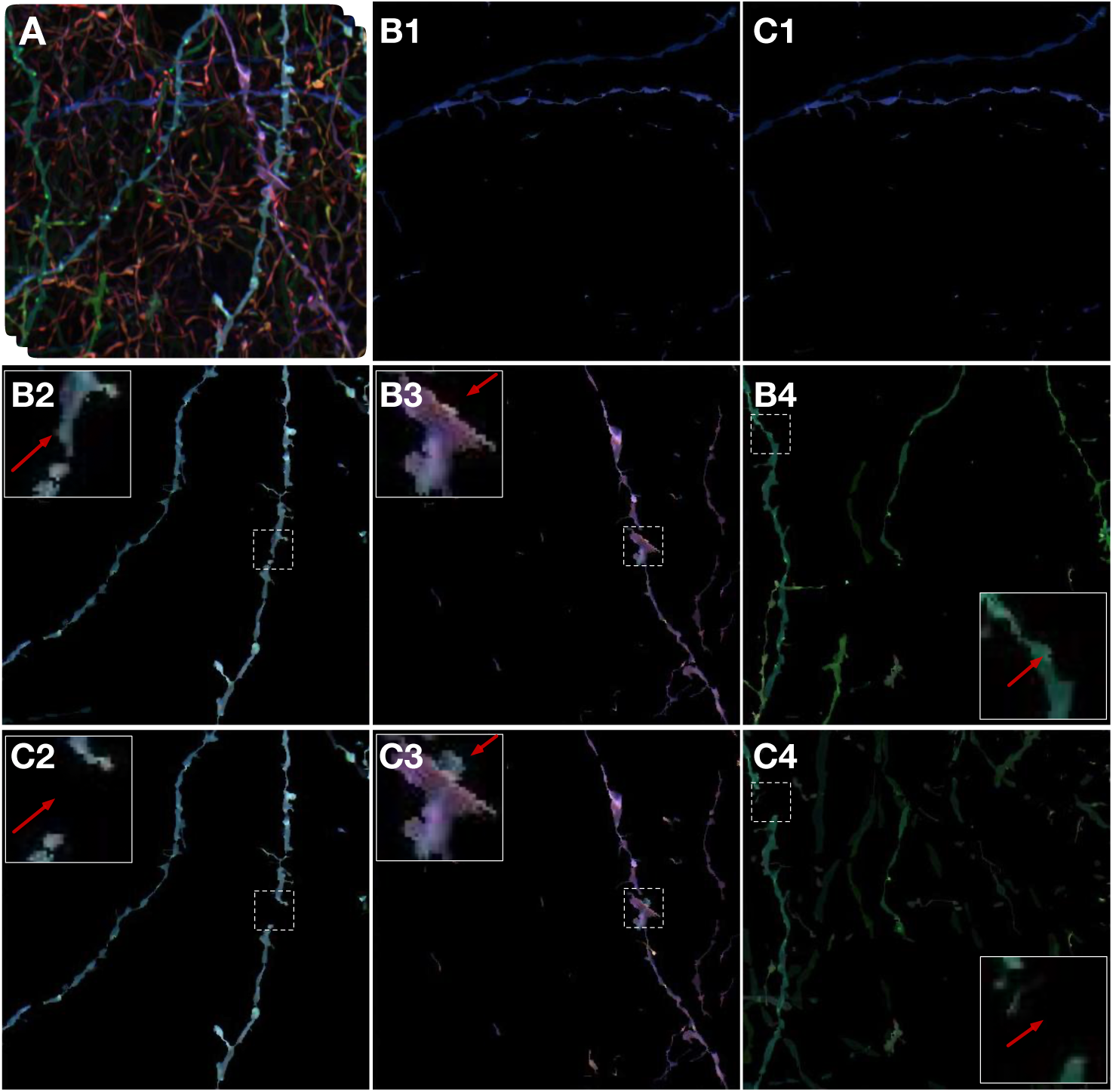
**A**, the input raw Brainbow image stack. **B1–4**, selected GMM segmentation results. **C1–4**, selected kernel *k*-means segmentation results. Insets are provided to show detail. The complete segmentation result can is visualized in Supp. Figure 1 and the tracings are available in Supp. Figures 2, 3. All the images are using maximum intensity projections along the *z*-axis and are rotated for better visualization.

### 2.4 Neuron Tracing

Despite the improvements in image segmentation, we still observe fragmented structures (Figure 2**D**) among the inferred neurons. To generate a compact tracing, we propose a method to merge fragmented topologies into one continuous tree. The fragmented neurites can be caused by the supervoxelization process and occlusions by the other neurons in the raw Brainbow data; the former problem has been eased by the implementation of GMM, but the latter persists. Thus, we develop a graph-based method to “bridge” the broken links within the same neurite. We first skeletonize the segmentation and use the resulting skeleton in the following operations.

#### Definition 1.

*Given a skeleton represented by a set of points, using a 27-point stencil, we define core points with at least 3 neighbors as branch points, and core points with only 1 neighbor as end points. This is demonstrated in Figure 3*.

#### Algorithm 1 Skeleton graph construction and linkage bridging

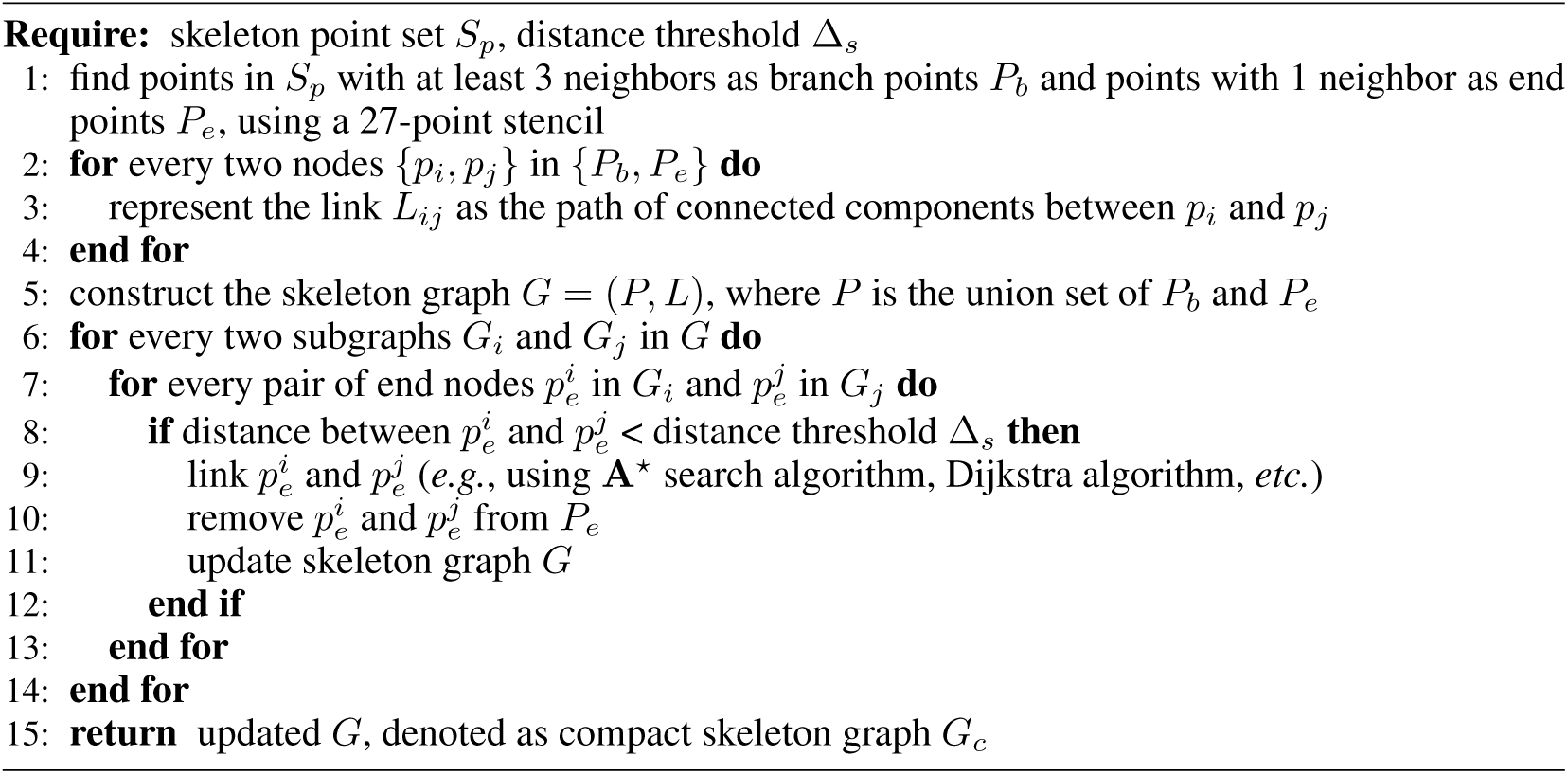

#### Skeleton graph construction

We represent branch points *P*_*b*_ and end points *P*_*e*_ as the nodes (i.e., branch and end nodes) in the graph, where the edges or links *L* between every two connected nodes are coded by the paths of the connected components between two nodes (Figure 3**B**). Note that, in this way, the points with 2 neighbors in the set of skeleton points can be well-represented by the links, and thus there is no need to represent these points as nodes. Abandoning the points with two neighbors also reduces the computation and improves efficiency. The skeleton graph is configured as *G* = (*P, L*), where *P* is the union set of branch nodes *P*_*b*_ and end nodes *P*_*e*_.

#### Linkage bridging

Broken linkages occur when two end points are incorrectly formed on opposite ends of a fragmented segmentation. Here, our way of constructing the graph is well-suited for bridging these broken connections. First, we extract subgraphs based on the connectivity of all the nodes. Next, we examine every pair of end nodes within two different subgraphs. When the distance, in our case a Euclidean distance, is less than threshold Δ_*s*_, we link the two end nodes, and thus we can obtain a more compact graph. The process is iterated until all pairs of subgraphs have been examined. A detailed description of this process is given in Algorithm 1, and we also show an alternative implementation using *k*-d tree in Supp. Algorithm 1.

#### Trace generation

In order to perform quantitative analysis from generated tracing, we developed a technique to generate SWC tracing files (Stockley et al., 1993). The SWC file format is used broadly by the neuroinformatics community and employed as the transaction format of various Fiji (Schindelin et al., 2012) plugins for calculating neuromorphology features. Our technique makes full use of the properties of *G*_*c*_. In general, a neurite starts from an end node and stops at an end node and the path between two connected end nodes can be interpreted as part of the neurite. In practice, we try to find the shortest paths from the seed node 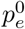 to following end nodes 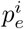 in the same subgraph, such that the union set of these found shortest paths can be interpreted as the final tracing result *T* (Tracing) which is of the form

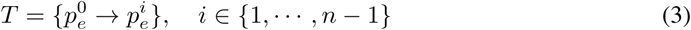

where *n* is the number of end nodes in the compact skeleton graph *G*_*c*_. We note that 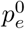 can be randomly selected from any end node, determined using a set of criteria (e.g., the end node furthest to the left of the image), or manually input. For our comparisons, we manually selected the end nodes to be consistent with our human annotation or chose 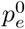 to be the end node with the smallest coordinate in unannotated data.

### 2.5 Trace Comparison

To quantify differences between different neuron tracings, we make use of the DIADEM metric (Gillette et al., 2011), which was derived as a way of objectively comparing the topologies of different neuron reconstructions. We used the published evaluation code, which generates a maximum value of 1 for perfectly-matching reconstruction, and 0 for a reconstruction which has no matched nodes to the gold standard. It is worth noting, however, that the DIADEM metric heavily weighs the branching topology of the neurite so, occasionally, visually-similar neurons will result in poor DIADEM scores. When running the DIADEM code, we used the parameter default preset ‘1’, while additionally lessening the X-Y-Z node-matching distance to 10 pixels, to account for the diameter of large-caliber neurites in the test image.

## 3 Datasets

We generated two test images for validation of our method in this report. Expansion microscopy (Shen et al., 2020) was applied to physically expand brain tissue by 4× from a Brainbow-labeled PV/Som-Cre mouse. A 3-channel image was collected, followed by manual channel alignment and histogram matched using Fiji (Schindelin et al., 2012). The resulting image was then cropped to 364 × 372 × 169 voxels, representing an effective voxel size of 75 nm × 75 nm × 175 nm and a physical volume of 27.3 µm × 27.9 µm × 29.6 µm, to form a manageable test case. This image is used in Figures 2, 4, 5, and 7.

**Figure 5:**
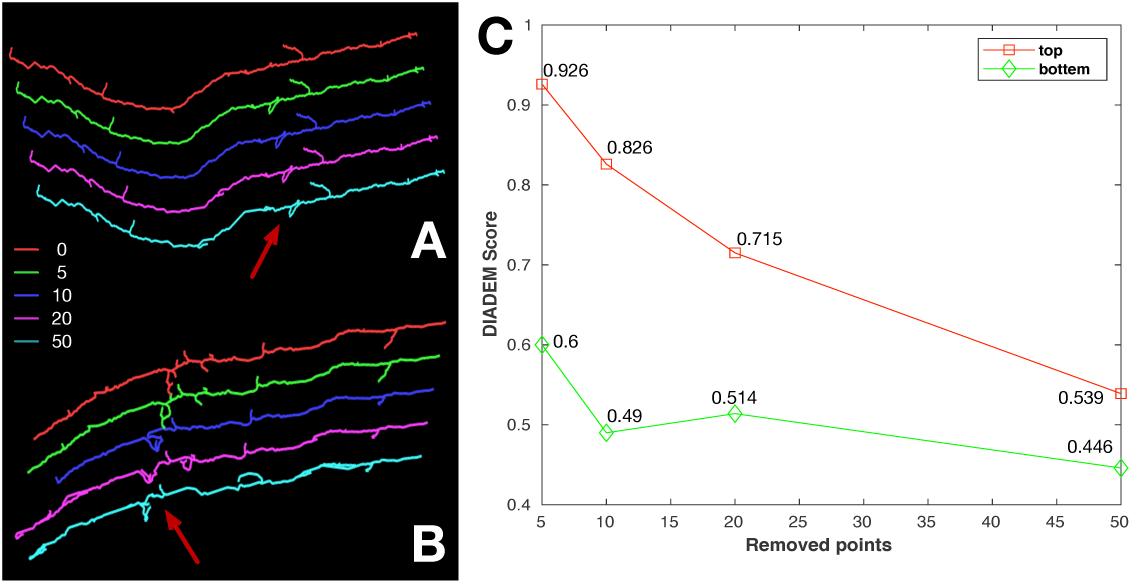
The study on robustness of our proposed method to loss of image information. **A** and **B**, comparisons on the tracing result of the left and right neurons found in Figure 4**B2. C**, the DI-ADEM quality score of each setting for **A** and **B**. Tracings have been rotated and rescaled to fit plotting area.

**Figure 6:**
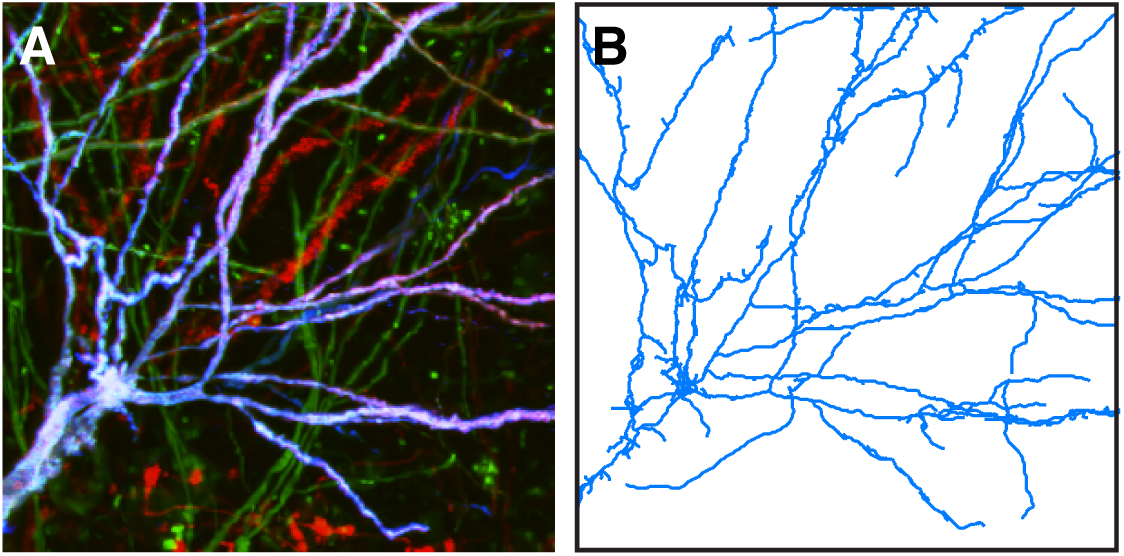
**A**, the projection of test image with interweaving neural branches along the *z*-axis. **B**, the tracing result using the proposed method for the purple neuron in the test image. Both images are maximum projections along the *z*-axis. Additional segmentation results are presented in Supp. Figure 4.

**Figure 7:**
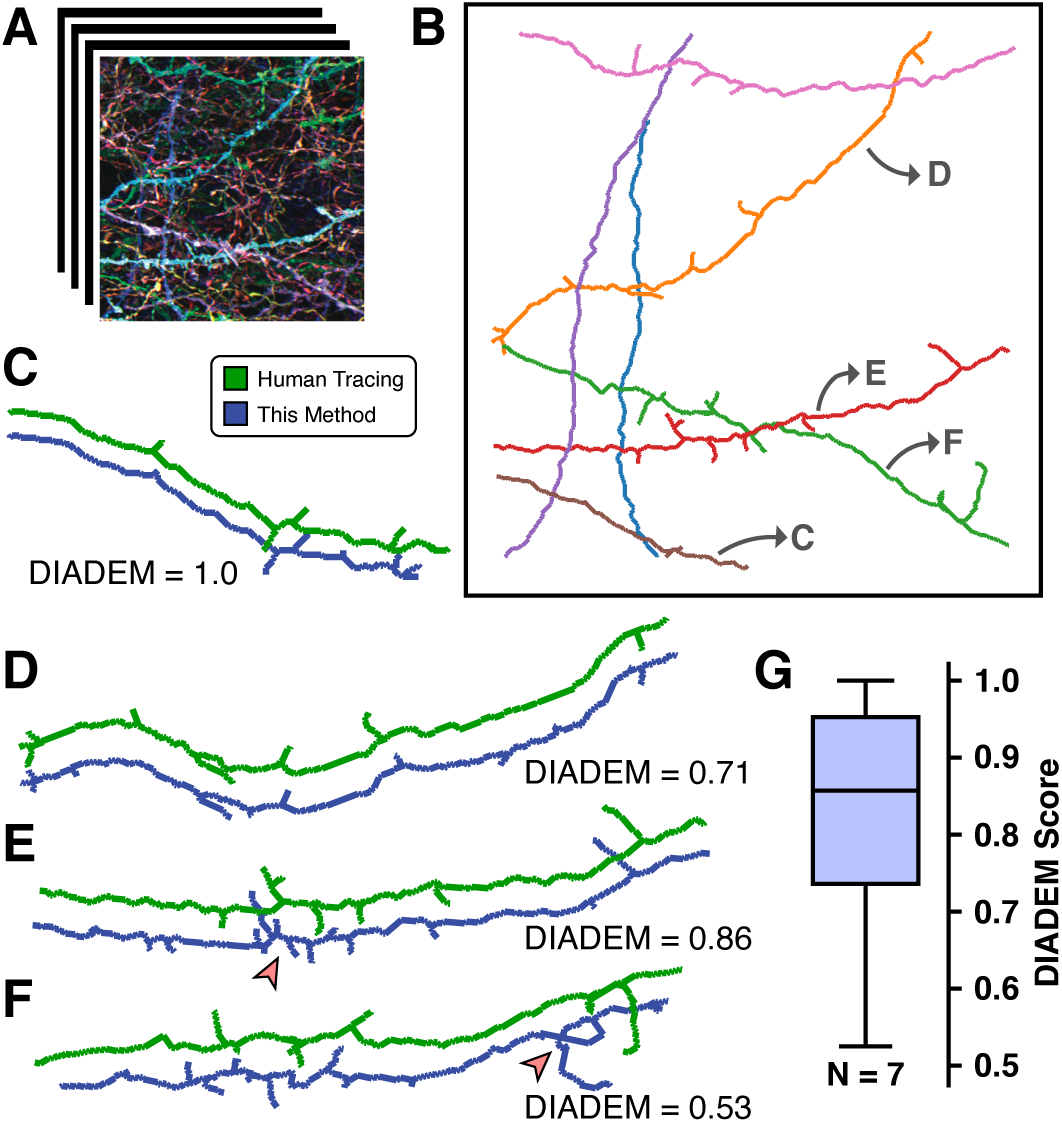
**A**, an overview *z*-projection of the test image, for reference. **B**, high caliber neurons (N = 7) were automatically reconstructed and their structures are visualized as *z*-projections of the resultant tracing files. **C–F**, several example neurons from **B** are shown next to manual human tracing. The DIADEM quality score for each pair is shown below each panel. Red arrows are discussed within the text. Note in **C** that slight differences exist between the two reconstructions, however, because they are smaller than the size limit for the algorithm, it reports a complete reconstruction. Neurons have been rotated and rescaled to fit the plotting area. **G**, the DIADEM scores of all 7 neurons in **B** (*µ* = 0.82; Range = [0.53, 1.0]).

We collected a second image (Figure 6) by injecting the Brainbow viral reporter into the hippocampal CA1 region of a POMC-Cre reporter mouse. 3-channel imaging was performed as above, without the use of sample expansion. This image was cropped to 300 × 300 × 300 voxels to encompass the branches of approximately a single neuron for demonstration purposes, and represents a voxel size of 0.42 µm × 0.42 µm × 1.00 µm and a physical volume of 126 µm × 126 µm × 300 µm.

## 4 Results

We first compare our neuron segmentation method (Methods 2.3) against the state-of-the-art method of Sümbül et al. by applying it to the first Brainbow image described above (Figure 4). Other than the number of clusters, the remaining parameters (described in Supp. Text) are held constant between the two methods, to ensure that differences observed are the result of algorithm changes. Overall, compared with the previous method, more continuous neuron processes are easily observed using GMM clustering. Channels which are well spectrally separated in the image result segmentations which are equivalent between the two methods (e.g., Figure 4**B1** and **C1**). When neurons of different color intersect, we find that our method results in fewer “extra” voxels being segmented to the wrong channel (Figure 4**B3** and **C3**; see arrows in insets). Finally, our method preserves continuous structures better, as observed by less degree of fragmentation in regions where calibers of the neurites vary (Figure 4**B2,4** and **C2,4**). These improvements result in segmentations that are more coherent and is crucial to accurate downstream automated data analyses.

Next, we implemented a neuron tracing routine to generate skeleton-like tree structure as neuron morphology presentation (Methods 2.4). To determine if our trace-generation method was robust to information loss introduced by segmentation errors, we performed a data loss simulation (Figure 5). Briefly, we calculated the skeletons for the two neurites found in Figure 4**B2**, after the random removal of between 0 and 50 points out of each 100, resulting in fragmented skeletons (Figure 5**A, B**). Upon evaluation, we find that both reconstructions are visually robust to large amounts of data loss, however, several loop structures are formed due to the loss of connectivity in dense regions (red arrows). The artificially-fragmented skeletons were then compared against the non-removal control using the DIADEM metric (Gillette et al., 2011) (Figure 5**C**). The quantification of the top neurite, as indicated by the DIADEM scores close to 1 indicates that there our linkage bridging method is robust against the loss of data in this case. We observe a similar trend in accuracy loss in the bottom neurite, however, the loop structures formed in error cause the DIADEM metric to be lower. Together, these results suggest that the qualitative structure of tracing is highly robust to the loss of data.

Then, we evaluated our method on a more complexly-branched sample as shown in Figure 6. Along with large color variance in the same neuron, the interweaving neural structure of this image can be hard even for human annotation. Figure 6**B** shows the result of bridging the fragmented segmentation into a compact tracing. We observe that even for the neurites with low intensity (Supp. Figure 4**B**, red arrows), our proposed can trace the neurites and add them into the reconstructed neuron tree. It is apparent that several branches are oversegmented, as a result of segmentation cross-talk with the green channel. This behavior can be tuned by parameter choice, however, we found that neuron “pruning” requires less human intervention time when proofreading a tracing as compared to adding missed branches.

Finally, as a test of the accuracy of our algorithm and applicability for large-scale neural circuit reconstruction experiments, we generated 7 neuron tracings from our test image (Figures 7**A, B**) and also reconstructed the same neurons by manual tracing. The automatically-generated tracings agree well with the human “gold standard” results (Figures 7**C–G**), with an average DIADEM score of 0.82. There are several features to be noted within these reconstructions: First, we find that there are some features which are reconstructed by the proposed method are not annotated by our human tracing (e.g., Figure 7**E**, arrow). Upon manual inspection, some of these small features represent spines that are difficult to resolve in the image. Additionally, one outlier reconstruction (Figure 7**F**) performed poorly, due to a small loop introduced by a nearby similarly-colored neurite (arrow). Overall, this experiment suggests the ability to perform large, automated reconstructions of Brainbow-labeled neurons with accuracy comparable to human annotation.

## 5 Discussion

In this paper, we present a method that enables the efficient generation of neuron structural traces from densely labeled multispectral Brainbow images. Specifically, our use of GMM clustering, as well as a graph-theoretic method for neuron trace repair, prevent fragmentation errors which result from the application of previous methods. We show by comparison to the human annotation of the same images that our method is robust and efficient while introducing minimal errors.

We hope that this work will find application with the many worldwide efforts to create whole-organism neural maps. Human proofreading time in these experiments can be astronomical, so improving automation has the potential to accelerate science by increasing its efficiency. Further, these developments may be relevant to similar tracing-extraction problems, such as retinal tracing.

### Broader Impact

Here, we present an unsupervised approach for neural tracing on densely labeled multispectral Brainbow images. Since it can relieve the high demand for human labor, our proposed method can help with the human annotations on Brainbow images which can potentially enables supervised or weakly-supervised approaches to learning brain circuits. Although approaches for reliably analyzing and quantifying Brainbow images remain in their infancy, we hope our work can draw the attention of researchers to this interesting field.

## Author Contributions

BD developed and implemented the algorithm. BD and LAW performed statistical analysis and drafted the manuscript, with input from all authors. DHR and FYS performed microscopy experiments. DC and YY conceptualized and oversaw the project.

## Acknowledgments

The authors thank Chaoxi Niu, Hao Tang, Jianfeng Cao, Ye Li, Jennifer Williams, and Nigel Michki for insightful discussions about this project.

DC received support by National Institutes of Health (NIH) 1UF1NS107659, 1R01MH110932, and 1RF1MH120005, and National Science Foundation NeuroNex-1707316. FYS received support by NIH 1F31NS11184701.

## References

James Baglama, Daniela Calvetti, and Lothar Reichel. Algorithm 827:irbleigs: A matlab program for computing a few eigenpairs of a large sparse hermitian matrix. ACM Transactions on mathematical software, 29(3):337–348, 2003.

Erhan Bas and Deniz Erdogmus. Piecewise linear cylinder models for 3-dimensional axon segmentation in brainbow imagery. In International symposium on biomedical imaging, pp. 1297–1300, 2010.

Dawen Cai, Kimberly B Cohen, Tuanlian Luo, Jeff W Lichtman, and Joshua R Sanes. Improved tools for the brainbow toolbox. Nature methods, 10(6):540–547, 2013.

Fei Chen, Paul W Tillberg, and Edward S Boyden. Expansion microscopy. Science, 347(6221): 543–548, 2015.

Inderjit S Dhillon, Yuqiang Guan, and Brian Kulis. Kernel k-means: spectral clustering and normalized cuts. In ACM SIGKDD International conference on knowledge discovery and data mining, pp. 551–556, 2004.

Todd A Gillette, Kerry M Brown, and Giorgio A Ascoli. The DIADEM metric: comparing multiple reconstructions of the same neuron. Neuroinformatics, 9(2-3):233, 2011.

James Gornet, Kannan Umadevi Venkataraju, Arun Narasimhan, Nicholas Turner, Kisuk Lee, H Sebastian Seung, Pavel Osten, and Uygar Sümbül. Reconstructing neuronal anatomy from whole-brain images. In International symposium on biomedical imaging, pp. 218–222, 2019.

Elizabeth MC Hillman, Venkatakaushik Voleti, Wenze Li, and Hang Yu. Light-sheet microscopy in neuroscience. Annual review of neuroscience, 42:295–313, 2019.

Wiebke Jahr, Benjamin Schmid, Christopher Schmied, Florian O Fahrbach, and Jan Huisken. Hyper-spectral light sheet microscopy. Nature communications, 6:7990, 2015.

Michal Januszewski, Jörgen Kornfeld, Peter H Li, Art Pope, Tim Blakely, Larry Lindsey, Jeremy Maitin-Shepard, Mike Tyka, Winfried Denk, and Viren Jain. High-precision automated reconstruction of neurons with flood-filling networks. Nature methods, 15(8):605–610, 2018.

Rui Li, Muye Zhu, Junning Li, Michael S Bienkowski, Nicholas N Foster, Hanpeng Xu, Tyler Ard, Ian Bowman, Changle Zhou, Matthew B Veldman, et al. Precise segmentation of densely interweaving neuron clusters using G-Cut. Nature communications, 10(1):1–12, 2019.

Ye Li, Logan A Walker, Yimeng Zhao, Erica M Edwards, Nigel S Michki, Hon Pong Jimmy Cheng, Marya Ghazzi, Tiffany Chen, Maggie Chen, Douglas H Roossien, and Dawen Cai. Bitbow: a digital format of brainbow enables highly efficient neuronal lineage tracing and morphology reconstruction in single brains. bioRxiv: 2020.04.07.030593, 2020.

Jean Livet, Tamily A Weissman, Hyuno Kang, Ryan W Draft, Ju Lu, Robyn A Bennis, Joshua R Sanes, and Jeff W Lichtman. Transgenic strategies for combinatorial expression of fluorescent proteins in the nervous system. Nature, 450(7166):56–62, 2007.

Matteo Maggioni, Vladimir Katkovnik, Karen Egiazarian, and Alessandro Foi. Nonlocal transform-domain filter for volumetric data denoising and reconstruction. IEEE Transactions on image processing, 22(1):119–133, 2012.

Fernand Meyer. Topographic distance and watershed lines. Signal processing, 38(1):113–125, 1994.

Darren Myatt, Tye Hadlington, Giorgio Ascoli, and Slawomir Nasuto. Neuromantic–from semi-manual to semi-automatic reconstruction of neuron morphology. Frontiers in neuroinformatics, 6:4, 2012.

Tingwei Quan, Hang Zhou, Jing Li, Shiwei Li, Anan Li, Yuxin Li, Xiaohua Lv, Qingming Luo, Hui Gong, and Shaoqun Zeng. Neurogps-tree: automatic reconstruction of large-scale neuronal populations with dense neurites. Nature methods, 13(1):51, 2016.

Douglas H Roossien, Benjamin V Sadis, Yan Yan, John M Webb, Lia Y Min, Aslan S Dizaji, Luke J Bogart, Cristina Mazuski, Robert S Huth, Johanna S Stecher, et al. Multispectral tracing in densely labeled mouse brain with ntracer. Bioinformatics, 35(18):3544–3546, 2019.

János Schanda. Colorimetry: understanding the CIE system. John Wiley & Sons, 2007.

Johannes Schindelin, Ignacio Arganda-Carreras, Erwin Frise, Verena Kaynig, Mark Longair, Tobias Pietzsch, Stephan Preibisch, Curtis Rueden, Stephan Saalfeld, Benjamin Schmid, et al. Fiji: an open-source platform for biological-image analysis. Nature methods, 9(7):676–682, 2012.

Hao-Chiang Shao, Wei-Yun Cheng, Yung-Chang Chen, and Wen-Liang Hwang. Colored multi-neuron image processing for segmenting and tracing neural circuits. In International conference on image processing, pp. 2025–2028, 2012.

Fred Y Shen, Margaret M Harrington, Logan A Walker, Hon Pong Jimmy Cheng, Edward S Boyden, and Dawen Cai. Light microscopy based approach for mapping connectivity with molecular specificity. bioRxiv: 2020.02.24.963538, 2020.

Colin JR Sheppard, Min Gu, and Maitreyee Roy. Signal-to-noise ratio in confocal microscope systems. Journal of Microscopy, 168(3):209–218, 1992.

EW Stockley, HM Cole, AD Brown, and HV Wheal. A system for quantitative morphological measurement and electrotonic modelling of neurons: three-dimensional reconstruction. Journal of Neuroscience methods, 47(1-2):39–51, 1993.

Uygar Sümbül, Douglas Roossien, Dawen Cai, Fei Chen, Nicholas Barry, John P Cunningham, Edward Boyden, and Liam Paninski. Automated scalable segmentation of neurons from multispectral images. In Advances in neural information processing systems, pp. 1912–1920, 2016.

Paul W Tillberg, Fei Chen, Kiryl D Piatkevich, Yongxin Zhao, Chih-Chieh Jay Yu, Brian P English, Linyi Gao, Anthony Martorell, Ho-Jun Suk, Fumiaki Yoshida, et al. Protein-retention expansion microscopy of cells and tissues labeled using standard fluorescent proteins and antibodies. Nature biotechnology, 34(9):987–992, 2016.

Tung-Yu Wu, Hung-Hui Juan, Henry Horng-Shing Lu, and Ann-Shyn Chiang. A crosstalk tolerated neural segmentation methodology for brainbow images. In International symposium on applied sciences in biomedical and communication Technologies, pp. 1–5, 2011.

Hang Xiao and Hanchuan Peng. App2: automatic tracing of 3d neuron morphology based on hierarchical pruning of a gray-weighted image distance-tree. Bioinformatics, 29(11):1448–1454, 2013.

Jian Yang, Ming Hao, Xiaoyang Liu, Zhijiang Wan, Ning Zhong, and Hanchuan Peng. FMST: an automatic neuron tracing method based on fast marching and minimum spanning tree. Neuroinformatics, 17(2):185–196, 2019.

Hongkui Zeng and Joshua R Sanes. Neuronal cell-type classification: challenges, opportunities and the path forward. Nature reviews neuroscience, 18(9):530–546, 2017.

