## Supplemental Materials for "Unsupervised Neural Tracing in Densely Labeled Multispectral Brainbow Images"

**Bin Duan**[1] **Logan A Walker**[2] **Douglas H Roossien**[3] **Fred Y Shen**[4]
**Dawen Cai**[*2,4,5] **Yan Yan**[*1]
[1]Department of Computer Science, Texas State University
[2]Biophysics, School of Literature, Science, and the Arts, University of Michigan
[3]Department of Biology, Ball State University
[4]Neuroscience Graduate Program, University of Michigan Medical School
[5]Department of Cell and Developmental Biology, University of Michigan Medical School

### Parameter Setting & Algorithm

**Noise standard deviation for denoising**: $\sigma = 1/8$ when the intensity is normalized to $[0, 1]$ for individual channels.

**Watershed transform**: 0.006 with 26-neighborhood. (By observation, the parameter which can range from 0.006 to 0.01 makes little difference on the quality of supervoxel.)

**LUV color features**: for images with more than 3 channels, for every normalized color triplet, we calculate the feature of in the LUV space. After concatenating all the LUV representations, we utilize principal components analysis to select the top C representations, where C is equal to the image channels.

**Edge construction:** spatial distance threshold $\delta_s = 50$, and color dissimilarity threshold $\delta_c = 0.5$. The spatial distance between two supervoxels is $min_{v \in V, u \in U} D(v, u)$, where $V$ and $U$ are the voxel sets of the two supervoxels and $D(v, u)$ is the Euclidean distance between voxels $v$ and $u$.

**Parameter of the Gaussian kernel**: $\gamma = 0.002$ for the weight decay on the edges.

**Feature dimensions**: $\mathbf{X} \in \mathbb{R}^{N \times d}$, where $N$ is the number of nodes (supervoxels), and $d$ is the numbers of top-$d$ largest eigenvalues. During experiments, it is setted around the number of the clusters.

**GMM clustering**: We adapt a variational approach (Chen, 2020) to perform GMM clustering. It allows rough parameter to get the final clustering result. That is, we set a number that is bigger than the actual number of the clusters. GMM will automatically find the optimal cluster number to converge. Specifically, we set 10 to Figure 4 and 5 to Figure 6 in the main text.

**Skeletonization**: While tracing is much more common in the study of connectomics and neuroscience, in order to produce lines by given the segmentation, we adapt some morphological operations (Kollmannsberger et al., 2017) to do the skeletonization on the segmentation, including erosion and dilation, *etc.*.

**27-point stencil**: end point with only 1 neighbor in the skeleton point set, and branch point with at least 3 neighbors in the skeleton point set.

**Skeleton graph construction**: we interpret each branch point or end point as node in the skeleton graph while the path between every two connected nodes are following the connected components in the skeleton point set using 27-point stencil.

**Linkage Bridging**: the linkage threshold $\Delta_s$ are 18 for Figure 4 and 8 for Figure 6 in main text. Here, we show one alternative implementation using $k$-d tree structures of linkage bridging which

is faster when the segmentation is heavily fragmented. Note that, heavily fragmented segmentation is hard to be reconstructed, therefore, more spurious link can be generated. We try to include more biological constraints in our future work. The alternative algorithm is depicted in Supplementary Algorithm 1.

**Supplementary Algorithm 1** Alternative linkage bridging

**Require:** skeleton point set $S_p$, distance threshold $\Delta_s$

find points in $S_p$ with at least 3 neighbors as branch points $P_b$ and points with 1 neighbor as end points $P_e$, using a 27-point stencil
**for** every two nodes $\{p_i, p_j\}$ in $\{P_b, P_e\}$ **do**
    represent the link $L_{ij}$ as the path of connected components between $p_i$ and $p_j$
**end for**
construct the skeleton graph $G = (P, L)$, where $P$ is the union set of $P_b$ and $P_e$.
**for** every end node $p_e^i$ **do**
    find the nearest end node $p_e^j$ using $k$-d tree structures
    **if** no connected path between $p_e^i$ and $p_e^j$ **then**
        **if** distance between $p_e^i$ and $p_e^j$ < distance threshold $\Delta_s$ **then**
            link $p_e^i$ and $p_e^j$ (*e.g.*, using $\mathbf{A}^\star$ search algorithm, Dijkstra algorithm, *etc.*)
            remove $p_e^i$ and $p_e^j$ from $P_e$
            update skeleton graph $G$
        **end if**
    **end if**
**end for**
**return** updated $G$, denoted as compact skeleton graph $G_c$

**Trace generation**: generating the tracing result enables post-processing in other softwares (Schindelin et al., 2012; Longair et al., 2011; Peng et al., 2010). The generation process is depicted in Supplementary Algorithm 2.

**Supplementary Algorithm 2** Trace generation

**Require:** compact skeleton graph $G_c$

list all end nodes $P_e$
set the seed end node $p_e^0$ automatically or manually
**for** every nodes $p_e^i$ in $P_e$ other than $p_e^0$ **do**
    starting from $p_e^0$, and following the connected path, find the shortest path $T_i = p_e^0 \to p_e^i$ between $p_e^i$ and $p_e^0$
    $currentpoint \leftarrow p_e^0$
    **while** $T_i$ has points **do**
        mark $currentpoint$ as parent node of the next point $t_i^j$ ($t_i^j$ starts right after $p_e^0$)
        remove $currentpoint$ from $T_i$
        $currentpoint \leftarrow t_i^j$
    **end while**
**end for**
**return** write SWC files

### Additional Results

**Reconstructed neural tracing.** We show the reconstructed neural tree topology for Figure 4**A** in main text in Supplementary Figure 2.

**More subtle neural process tracing and failures.** While Supplementary Figure 2 shows the performance of our method on high caliber neural process, we try to explore the proposed method on more subtle neural process — high caliber neurons. The details are showed in Supplementary Figure 3. Note that even for human, the subtle neural process is hard to be identified, while limited by both the segmentation result and the intricate neural tree structure, the tracing result seems worse than the main one. However, we can still observe some traced processes.

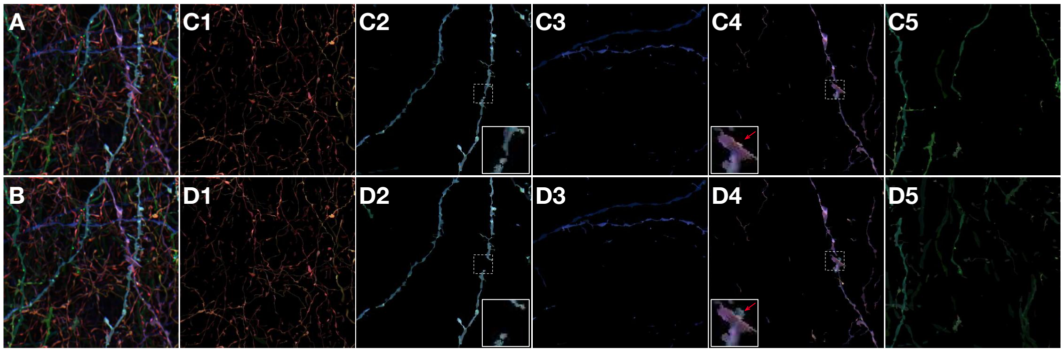


Supplementary Figure 1: The complete segmentation result of Brainbow image used in main text (Figure 4). **A** and **B** is the same input Brainbow image, **C1–C5**, segmentation result of GMM, **D1–D5**, segmentation result of kernel $k$-means. All the images are using maximum intensity projections along the $z$-axis.

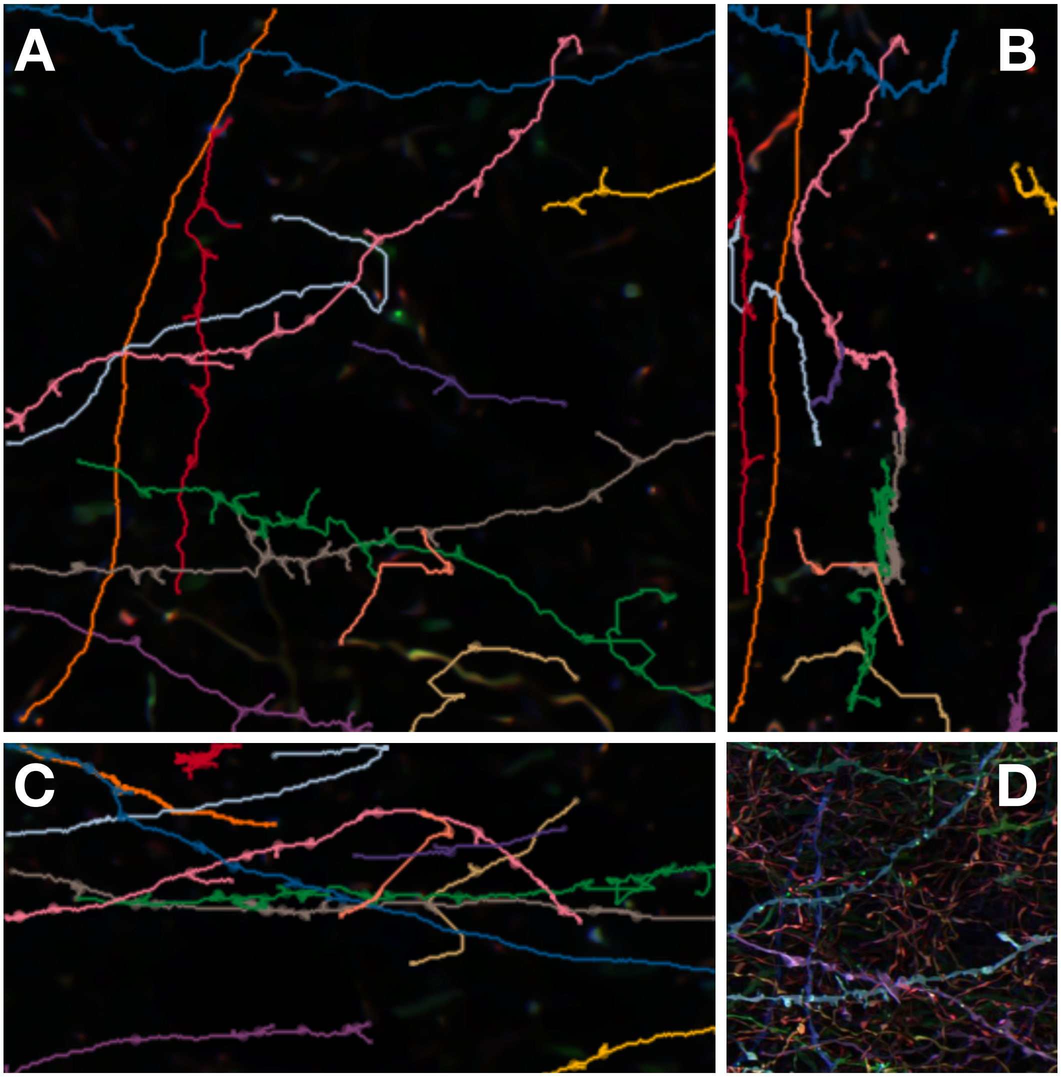


Supplementary Figure 2: **A**, XY view, **B**, XZ view, **C**, YZ view of the tracing. **D**, maximum intensity projection of the raw image in Figure 4 in main text. We can notice that the main process is compactly traced. All tracing results are pseudo-colored.

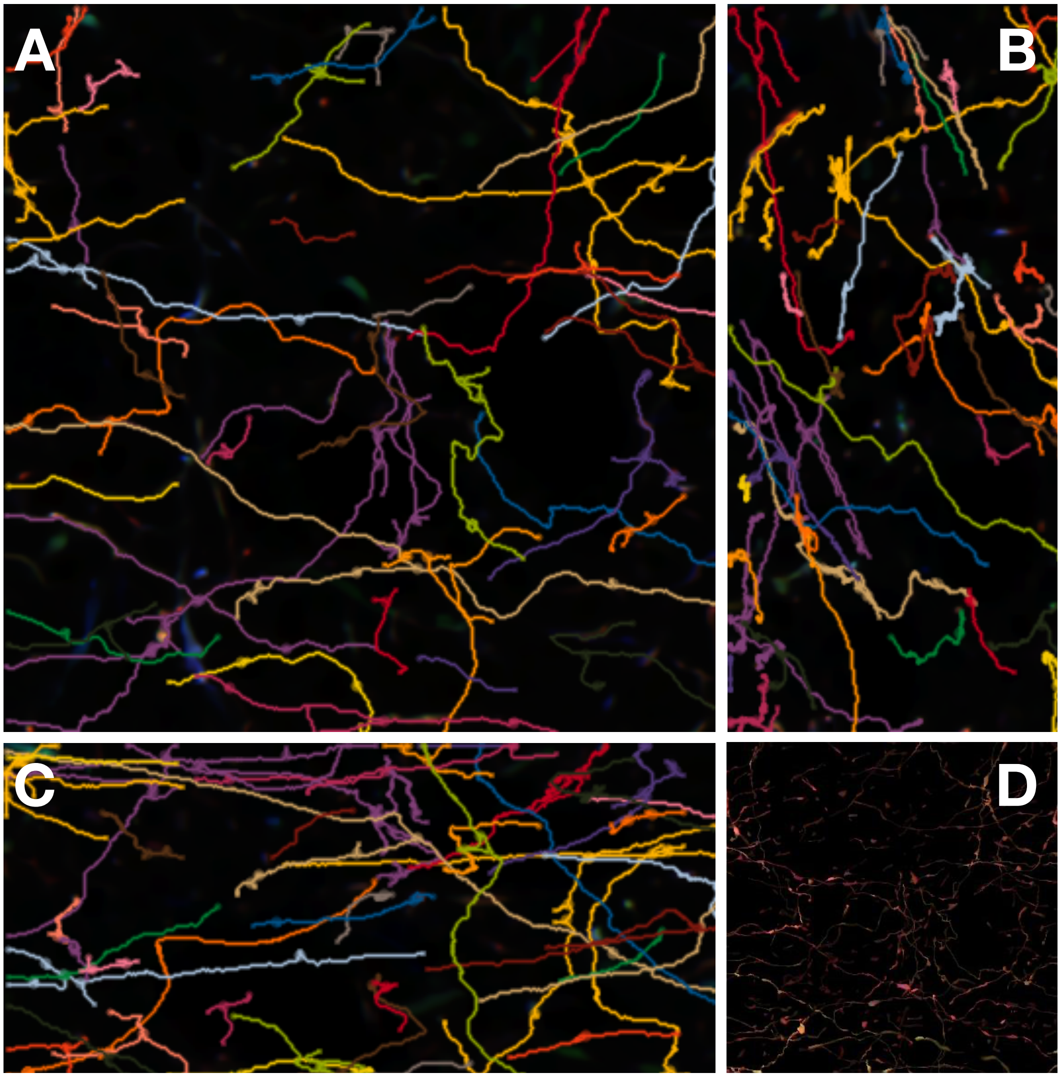


Supplementary Figure 3: **A**, XY view, **B**, XZ view, **C**, YZ view of the tracing. **D**, maximum intensity projection of the segmentation of the subtle neural process in Figure 4 in main text.

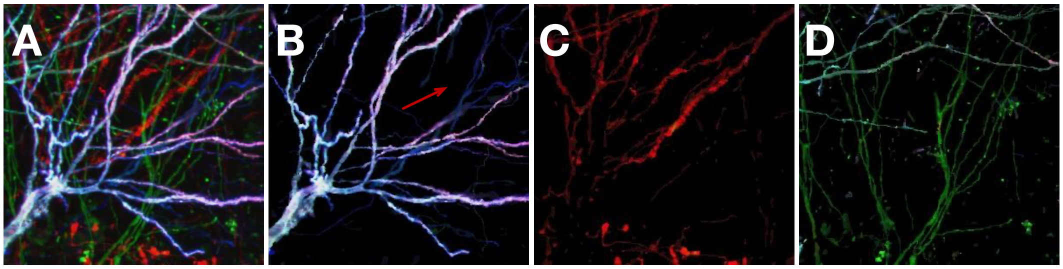


Supplementary Figure 4: The entire segmentation result of Brainbow image used in main text (Figure 6). **A**, the input Brainbow image, **B–D**, segmentation result. All the images are using maximum intensity projections along the $z$-axis.

### References

Mo Chen. Variational bayesian inference for gaussian mixture model. `https://www.github.com/PRML/PRMLT`, 2020.

Philip Kollmannsberger, Michael Kerschnitzki, Felix Repp, Wolfgang Wagermaier, Richard Weinkamer, and Peter Fratzl. The small world of osteocytes: connectomics of the lacuno-canalicular network in bone. *New Journal of Physics*, 19(7):073019, 2017.

Mark H Longair, Dean A Baker, and J Douglas Armstrong. Simple neurite tracer: open source software for reconstruction, visualization and analysis of neuronal processes. *Bioinformatics*, 27 (17):2453–2454, 2011.

Hanchuan Peng, Zongcai Ruan, Fuhui Long, Julie H Simpson, and Eugene W Myers. V3d enables real-time 3d visualization and quantitative analysis of large-scale biological image data sets. *Nature biotechnology*, 28(4):348–353, 2010.

Johannes Schindelin, Ignacio Arganda-Carreras, Erwin Frise, Verena Kaynig, Mark Longair, Tobias Pietzsch, Stephan Preibisch, Curtis Rueden, Stephan Saalfeld, Benjamin Schmid, et al. Fiji: an open-source platform for biological-image analysis. *Nature methods*, 9(7):676–682, 2012.
